# Coupling between tolerance and resistance for two related *Eimeria* parasite species

**DOI:** 10.1101/2020.01.24.918144

**Authors:** Alice Balard, Víctor Hugo Jarquín-Díaz, Jenny Jost, Vivian Mittné, Francisca Böhning, Ľudovít Ďureje, Jaroslav Piálek, Emanuel Heitlinger

## Abstract

Resistance (host capacity to reduce parasite burden) and tolerance (host capacity to reduce impact on its health for a given parasite burden) manifest two different lines of defence. Tolerance can be independent from resistance, traded-off against it, or the two can be positively correlated because of redundancy in underlying (immune) processes. We here tested whether this coupling between tolerance and resistance could differ upon infection with closely related parasite species. We tested this in experimental infections with two parasite species of the genus *Eimeria*. We measured proxies for resistance (the (inverse of) number of parasite transmission stages (oocysts) per gram of feces at the day of maximal shedding) and tolerance (the slope of maximum relative weight loss compared to day of infection on number of oocysts per gram of feces at the day of maximal shedding for each host strain) in four inbred mouse strains and four groups of F1 hybrids belonging to two mouse subspecies, *Mus musculus domesticus* and *M. m. musculus*. We found a negative correlation between resistance and tolerance against *E. falciformis*, while the two are uncoupled against *E. ferrisi*. We conclude that resistance and tolerance against the first parasite species might be traded off, but evolve more independently in different mouse genotypes against the latter. We argue that evolution of the host immune defences can be studied largely irrespective of parasite isolates if resistance-tolerance coupling is absent or weak (*E. ferrisi*) but host-parasite coevolution is more likely observable and best studied in a system with negatively correlated tolerance and resistance (*E. falciformis*).

## Introduction

Host defence mechanisms evolve to alleviate the detrimental effect of parasites. They can be categorised into two components: resistance and tolerance (Råberg et al. 2009). Resistance is the ability of a host to reduce parasite burden, resulting from defence against parasite infection or proliferation early after infection (Schmid-Hempel 2013). The negative effect of resistance on parasite fitness can lead to antagonistic coevolution. According to theoretical models, fluctuating host and parasite genotypes arise, and balancing selection maintains resistance alleles polymorphic (Boots et al. 2008; Roy & Kirchner 2000). Resistance has been the classical “catch all” measure for host-parasite systems, but recently it has been shown to be incomplete, especially with respect to potential fitness effects on the host (Kutzer & Armitage 2016; Råberg et al. 2009).

Disease tolerance (not to be confused from “immunological tolerance”, unresponsiveness to self antigens; Medzhitov et al. 2012) is the ability of the host to limit the impact of parasite on its fitness (Råberg et al. 2009; Vale & Little 2012; Kutzer & Armitage 2016). By potentially providing a longer-living niche, this defence mechanism improves, or at least does not deteriorate, the fitness of the parasite. Tolerance alleles are thus predicted by theoretical models to evolve to fixation due to positive feedback loops (Boots et al. 2008; Restif & Koella 2004; Roy & Kirchner 2000). From a mechanistic perspective tolerance alleviates direct or indirect damage (e.g. excessive immune response underlying resistance against parasites, called immunopathology; Graham et al. 2005) caused by parasites (Råberg et al. 2009). Tolerance mechanisms include modulation of inflammatory response (Ayres & Schneider 2012), tissue repair (stress response, damage repair and cellular regeneration mechanisms; Soares et al. 2017), and compensation of parasite-induced damage by increase of reproductive effort (Baucom & Roode 2011). Even in the absence of parasite infection, the maintenance of tolerance mechanisms can be detrimental to other functions, ultimately affecting host fitness (Stowe et al. 2000; Råberg et al. 2009). The resulting costs of resistance and tolerance determine the optimal (steady state and infection inducible) extent of both immune defences (Sheldon & Verhulst 1996).

Resistance and tolerance can be positively associated if they involve the same metabolic pathway, as was shown in the plant model *Arabidopsis thaliana* in response against herbivory (Mesa et al. 2017). In animals, genetic association studies of resistance and tolerance of *Drosophila melanogaster* against the bacterium *Providencia rettgeri* have shown positively correlated genetic effects, as the same loci were associated with changes of both traits in the same direction (Howick & Lazzaro 2017).

Nevertheless, resistance and tolerance can also be genetically and physiologically independent, involving different proximate mechanisms. Lack of correlation between both defences was shown for example in monarch butterflies (*Danaus plexippus*) infected by the protozoan parasite *Ophryocystis elektroscirrha*. This study found genetic variation in resistance between butterflies families, but a fixed tolerance (Lefèvre et al. 2010). Similarly, no correlation could be found between resistance and tolerance for the fish *Leuciscus burdigalensis* in response to infection with its parasite *Tracheliastes polycolpus*. The authors explain the decoupling of both defences by the fact that, in this system, tolerance likely involves wound repair rather than immune regulation, making resistance and tolerance mechanisms independent (Mazé-Guilmo et al. 2014).

In other systems, resistance and tolerance have been found negatively correlated. For example, inbred laboratory mouse strains lose weight upon infection with *Plasmodium chabaudi*. The extent of this impact on host health is negatively correlated with the peak number of parasites found in the blood (Råberg et al 2007), meaning that mouse strains with higher resistance present lower tolerance. Similarly, infections of sea trout (*Salmo trutta trutta*) and Atlantic salmon (*Salmo salar*) with the trematode *Diplostomum pseudospathaceum* showed that resistance and tolerance were negatively correlated when assessing mean levels of both traits in different host populations (Klemme & Karvonen 2016). This is interpreted as a result of trade-off between resistance and tolerance (Sheldon & Verhulst 1996; Restif & Koella 2004; Råberg et al. 2009).

We have seen that depending on the system studied, resistance and tolerance can be (1) uncoupled (independent), (2) positively correlated (involving same genes and mechanisms), or (3) negatively correlated (traded-off). Theoretical models show that coupling between resistance and tolerance (or absence thereof) could depend not only on the host but also on the parasite (Carval & Ferriere 2010). Here we tested this hypothesis. More precisely, we asked whether there could be differences in the resistance-tolerance coupling upon infection of one host type with two closely related parasite species. To answer this question, we infected four inbred mouse strains and four groups of F1 hybrids representative of two house mouse subspecies, *M. m. domesticus* and *M. m. musculus*, with two parasite isolates representative of two naturally occurring parasite species, the protozoan parasites *Eimeria ferrisi* and *E. falciformis* (Jarquín-Díaz et al. 2019). *Eimeria* spp. are monoxenous parasites that expand asexually and reproduce sexually in intestinal epithelial cells, leading to malabsorption of nutrients, tissue damage and weight loss (Chapman et al. 2013). The evolutionary history of these different *Eimeria* species in the two house mouse subspecies is unknown and it is unclear whether subspecies-specific adaptation exists in one or the other. We tested if coupling between resistance and tolerance differs between both parasite species and discussed the implication for parasite-host coevolution.

## Material and methods

### 1. Parasite isolates

The three parasite isolates used in this study were isolated from feces of three different *M. m. domesticus/M. m. musculus* hybrid mice captured in Brandenburg, Germany, in 2016 (capture permit No. 2347/35/2014). The parasite isolates belong to both the most prevalent *Eimeria* species in this area, namely *E. ferrisi* (isolate Brandenburg64) and *E. falciformis* (isolate Brandenburg88)(Jarquín-Díaz et al. 2019). Isolate Brandenburg64 was isolated in a 92% *M. m. domesticus* individual (hybrid index (HI) = 0.08: Proportion of *M. m. musculus* alleles in a set of 14 diagnostic markers, see Balard et al. (2020)) and isolate Brandenburg88 in a 80% *M. m. domesticus* (HI=0.2). Pre-patency and the peak day of parasite shedding for these isolates were estimated during infection in NMRI laboratory mice (Al-khlifeh et al. 2019) which were also used for serial passaging of the isolates. Previous to the experiment, the isolates had been passaged respectively 3 and 4 times in NMRI laboratory mice. Parasite infective forms (oocysts) were recovered by flotation in saturated NaCl solution followed by washing and observation under light microscope (following the protocol described in Clerc et al. (2019)) and stored at room temperature in 1mL of 2% potassium dichromate for a maximum of 2 months before infection of the wild-derived mice. Oocysts were allowed to sporulate 10 days before infection in a water bath at 30°C.

### 2. Mouse groups

We used four wild-derived inbred mouse strains from which we generated four groups of F1 hybrids. Hybrids between *M. m. domesticus* and *M. m. musculus* are used in the present study solely to increase statistical power for comparisons among strains (such as resistance-tolerance correlations). In the future, analyses of a hybrid effect (Balard et al. 2020) could investigate tolerance and resistance employing a larger panel of such hybrid strains allowing statistical analysis of an outbreeding effect. Two parental strains represented *M. m. domesticus*: **SCHUNT** (Locality: Schweben, Hessen, Germany [N: 5°0 26’, E: 9° 36’] (Martincová et al. 2019)) and **STRA** (Locality: Straas, Bavaria, Germany [N: 50° 11’, E: 11° 46’] (Piálek et al. 2008), and two derived from *M. m. musculus*: **BUSNA** (Locality: Buškovice, Bohemia, Czech Republic [N: 5°0 14’, E: 1°3 22’] (Piálek et al. 2008)) and **PWD** (Locality: Kunratice, Bohemia, Czech Republic [N: 5°0 01’, E: 14 2°9’] (Gregorová and Forejt 2000)). These four strains were fully inbred, i.e. passing more than 20 generations of brother–sister mating. The four groups of F1 hybrids consisted of two intrasubspecific hybrids (**SCHUNTxSTRA** and **PWDxBUSNA**) and two intersubspecific hybrids (**STRAxBUSNA** and **SCHUNTxPWD**) (Figure 1). Age of the mice at the time of infection ranged between 5.6 and 21.4 weeks, with the mean for each eight mouse group ranging between 10.5 and 14.7 weeks. All mouse strains and F1 hybrids were obtained from the Institute of Vertebrate Biology of the Czech Academy of Sciences in Studenec (license number 61974/2017-MZE-17214; for further details on strains see https://housemice.cz/en).

**Figure 1.**
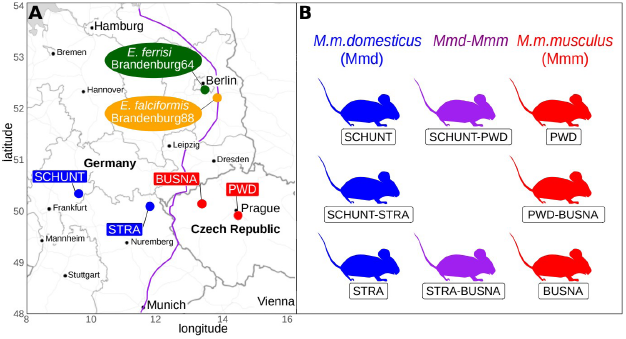
Parasite isolates and mouse wild-derived strains. (A) Map showing locations at which mice were collected for breeding of mouse strains and isolation of parasites. The purple line is an estimation of the center of the house mouse hybrid zone between *M. m. domesticus* and *M. m. musculus* based on sampling and genotyping of mice in this area (Balard et al. 2020; Ďureje et al. 2012; Macholán et al. 2019). (B) The eight mouse groups (parents and F1s) used in our experimental infections.

Parasites of the *Eimeria* genus are known to induce host immune protection against reinfection (Rose, Hesketh, and Wakelin 1992; Smith and Hayday 2000). To ensure that our mice were *Eimeria*-naive, mouse fecal samples were tested before infection for the presence of *Eimeria* spp. oocysts by flotation in saturated NaCl solution followed by washing and observation under light microscope.

### 3. Experimental infection

Mice were kept in individual cages during infection. Water and food (SNIFF, Rat/Mouse maintenance feed 10 mm) were provided *ad libitum* supplemented with 1 g of sunflower and barley seeds per day. Mice were orally infected with 150 sporulated oocysts of one *Eimeria* isolate suspended in 100*μ*l phosphate-buffer saline (PBS) and monitored daily until their sacrifice by cervical dislocation at time of regression of infection (reduction of oocyst output). Individuals presenting severe health deficiency and/or a weight loss approaching 18% relative to their starting weight were sacrificed earlier at defined humane end points (experiment license Reg. 0431/17). Weight was recorded and feces collected on a daily basis. Fecal pellets were collected every day from each individual cage and suspended in 2% potassium dichromate. Parasite oocysts were recovered using NaCl flotation (see above).

All individuals were negative for *Eimeria* at the beginning of our experiment (before infection of first batch, as described in the next paragraph). In total, 143 mice were infected. Mice were randomly allocated to experimental groups ensuring homogeneous distribution of ages and sexes between groups. Our experiments were conducted in four (partially overlapping) consecutive batches for logistical reasons. The first two batches were infected with *E. ferrisi* isolates (Brandenburg64), the third and fourth by one *E. ferrisi* isolate (Brandenburg64) and one *E. falciformis* isolate (Brandenburg88). Our experimental design is summarized in Table 1 (chronology of experimental batches can be scrutinized in Appendix 1).

**Table 1.** Infection experiment design.

| Host |  | Parasite |  |
| --- | --- | --- | --- |
| Mouse strains | Mouse subspecies | *E. ferrisi*<br>Brandenburg64 | *E. falciformis*<br>Brandenburg88 |
| SCHUNT | *F0 M.m.domesticus* | 14 (6M / 8F) | 6 (3M / 3F) |
| STRA | *F0 M.m.domesticus* | 15 (8M / 7F) | 7 (4M / 3F) |
| SCHUNTxSTRA | *F1 M.m.domesticus* | 6 (2M / 4F) | 8 (5M / 3F) |
| STRAxBUSNA | *F1 Hybrid* | 8 (5M / 3F) | 8 (3M / 5F) |
| SCHUNTxPWD | *F1 Hybrid* | 8 (3M / 5F) | 6 (4M / 2F) |
| PWDxBUSNA | *F1 M.m.musculus* | 9 (4M / 5F) | 7 (4M / 3F) |
| BUSNA | *F0 M.m.musculus* | 14 (8M / 6F) | 7 (3M / 4F) |
| PWD | *F0 M.m.musculus* | 13 (10M / 3F) | 7 (1M / 6F) |

Nematode infection is common in breeding facilities (Baker, 1998) and could interact with *Eimeria* (Clerc et al. 2019). We surveyed for their presence and nematode eggs (*Syphacia* sp. and *Aspiculuris* sp.) were observed in flotated feces of mice belonging to all genotypes before the experiment. Despite treatment of the first infection batch of mice (B1, 12 mice) with anthelminthics (Profender®, Bayer AG, Levekusen, Germany) following the protocol of Mehlhorn et al. (2005), nematodes were still detected with PCR (following the protocol of (Floyd et al. 2005)) in randomly sampled fecal samples a week later. We therefore decided not to treat mice of the following infection batches. Moreover, we observed *Eimeria* oocysts in the feces of 28 mice belonging to the last experimental batch (batch B4) at the day of infection, likely due to cross-contamination between batches. For following statistical analyses, we considered along with the full data set (N=143) a conservative data set in which cross-contaminated animals and animals treated by anthelminthic were removed (N=103). Results obtained on the conservative data set can be found in Appendix 2 and 3. Despite differences in significance due to a lower statistical power, the main conclusions of our analyses were consistent with those obtained on the main data set.

### 4. Statistical analyses

#### 4.1. Choice of proxies for resistance, impact of parasite on host and tolerance

As resistance is the capacity of a host to reduce its parasite burden, it is usually estimated by the inverse of infection intensity (Råberg et al. 2009). Pre-patency (the time to shedding of infectious stages, so called oocysts) is longer for *E. falciformis* (7 days) than for *E. ferrisi* (5 days) (Al-khlifeh et al. 2019). Therefore, as a proxy of (inverse of) resistance we used the number of oocysts per gram of feces (OPG) at the day of maximal shedding. Using the Spearman’s non-parametric rank correlation test, we found this measurement to be tightly correlated with the sum of oocysts shed throughout the experiment (Spearman’s *ρ*=0.93, N=168, P<0.001). Due to the aggregation characteristic of parasites (Shaw and Dobson 1995), the appropriate distribution for maximum number of OPG was found to be the negative binomial distribution. This was confirmed based on log likelihood, AIC criteria and goodness-of-fits plots (density, CDF, Q-Q, P-P plots; R packages MASS (Venables & Ripley 2002) and fitdistrplus (Delignette-Muller & Dutang 2015)). We confirmed the fit of our models by assessing the uniformity of the distribution of model residuals.

Both parasite species provoke inflammation, cellular infiltration, enteric lesions, diarrhea, and ultimately weight loss (Ankrom, Chobotar, and Ernst 1975; Ehret et al. 2017; Schito, Barta, and Chobotar 1996; Al-khlifeh et al. 2019). Therefore, the impact of parasites on host health was measured as the maximum relative weight loss compared to day 0 (body weight measured at the start of the experimental infection). For mice sacrificed at humane end points before the end of the experiment, the last weight of the living animal was used. This weight (loss) can be expected to be a very conservative estimate for our analyses (rendering tolerance conservatively low for these animals, which might have lost more weight if not sacrificed).

Tolerance is usually defined as a reaction norm, i.e. the regression slope of host fitness (or health condition if that is the parameter of interest) on infection intensity per host genotype (Simms 2000; Råberg et al. 2009). Thus tolerance was assessed as the slope of maximum relative weight loss compared to day 0 on number of OPG at the day of maximal shedding, within each mouse group and for each parasite isolate. A steep slope indicates a low tolerance (high weight lost for a given parasite burden).

#### 4.2. Statistical comparison of resistance, impact on health and tolerance in *E. ferrisi* and *E. falciformis*

The comparison between *E. ferrisi* and *E. falciformis* was performed using respectively the isolates Brandenburg64 and Brandenburg88 with which we infected all our eight mouse groups (see Table 1). Maximum OPG and relative weight loss were modelled separately as a response of mouse group, parasite isolate and their interaction. We used a negative binomial generalised linear model for maximum OPG, and a linear model for relative weight loss. Tolerance was assessed by modelling relative weight loss as a response of maximum OPG interacting with mouse group, parasite isolate and the interaction of the two latter. As each mouse was controlled against itself at the start of the experiment, before losing weight or shedding parasites, we performed a linear regression with null intercept. To test the significance of the marginal contribution of each parameter to the full model, each parameter was removed from the full model, and the difference between full and reduced model was assessed using likelihood ratio tests (G).

For each of our models that showed a significant interaction term, we also asked within each parasite isolate if the response differed between mouse groups using likelihood ratio tests (G) as described above. In the case of a non-significant interaction term, we performed post-hoc tests corrected for multiple testing (Tukey Honest Significant Differences (HSD)) to compare within all pairwise comparisons between groups (parasite isolate-mouse strain).

Of note, four mice infected with *E. falciformis* isolate Brandenburg88 did not shed any oocysts as death occurred at or one day before the peak of oocysts shedding in other mice. For this reason, we modelled maximum OPG when mice infected with this parasite were included using a zero-inflated negative binomial (ZINB) generalised linear model, after verifying that it provided a better fit than the simple negative binomial based on log likelihood and AIC criteria.

#### 4.3. Test of coupling between resistance and tolerance

We tested coupling between resistance and tolerance for *E. ferrisi* and *E. falciformis* using the isolates Brandenburg64 and Brandenburg88 and our eight mouse groups. To test such coupling, one can assess the strength of correlation between measure of resistance and measure of tolerance (Råberg et al 2007). Of note, tolerance (in absolute value) is measured as the slope *α* of the linear regression of parasite load (x) on maximum relative weight loss (y) of equation y = *α* x + *β* (*α* being the slope and *β* the intercept, 0 in our case). Therefore, tolerance is expressed as *α* = y/x – *β*/x. As x and y/x are by definition not independent, testing the correlation between resistance and tolerance can lead to spurious correlation (Brett 2004). To alleviate the dangers of this statistical artifact, we additionally tested differences in resistance, impact on health and tolerance between mouse groups separately (as described before, see 4.2) and also the underlying correlation between mean parasite load (x) and mean relative weight loss (y). We use the terminology “coupling” (between resistance and tolerance) to describe genotype-level correlation between tolerance and resistance additionally supported by the absence of positive correlation between health-effect and resistance. Correlations were tested using Spearman’s rank correlation.

After testing the resistance-tolerance coupling separately in both parasites, we tested the statistical difference in the relationship between (1) health-effect and resistance and (2) tolerance and resistance in the two *Eimeria* species infections. To achieve this aim, we used the mean values predicted by our three models (see 4.2) for each eight mouse groups to perform first a linear regression of the mean predicted relative weight loss as a response of the mean predicted OPG, parasite isolate and their interaction, and second a linear regression of the mean predicted tolerance value as a response of the mean predicted OPG, parasite isolate and their interaction. The significance of the marginal contribution of each parameter to the full model was assessed by removing each parameter from the full model, and the difference between full and reduced model was assessed using likelihood ratio tests (G).

All analyses were performed using R version 3.5.2 (R Development Core Team 2013) (negative binomial: function glm.nb from R package MASS (Venables and Ripley 2002); ZIBN: function zeroinfl from R package pscl (Jackman 2020; Zeileis, Kleiber, and Jackman 2008); linear model: function lm from R core package stats; mean and 95% confidence intervals: function ggpredict from R package ggeffect (Lüdecke 2018)). Graphics were produced using the R package ggplot2 (Wickham 2016) and compiled using the free software inkscape (https://inkscape.org).

## Results

### 1. General

Parasites of all isolates successfully infected all mouse groups (at the exception of 5 individuals infected with the *E. falciformis* isolate Brandenburg88 that died or had to be sacrificed due to a strong weight loss before the peak of shedding for this parasite), meaning that no “qualitative infection resistance” (*sensu* (Gandon and Michalakis 2000)) was detected. For *E. ferrisi* isolate Brandenburg64, the pre-patent period was 5 days post infection (dpi) and the median day of maximal oocyst shedding was 6 dpi (standard deviation sd=0.9). The median day of maximum weight loss was 5 dpi for both isolates (sd=1.7). For *E. falciformis* isolate Brandenburg88 pre-patency was 7 dpi, median day of maximal shedding was 8 dpi (sd=1.3) and median day of maximal weight loss 9 dpi (sd=1.6)(Figure 2). Of note a considerable number of mice infected with this isolate (13 out of 56 = 23%) died or had to be sacrificed at humane end points less than 3 days after the oocysts shedding peak for the group, all belonging to *M. m. musculus* subspecies (PWD, BUSNA, or their F1 PWDxBUSNA; 5 died at dpi 8, 5 at dpi 9, 3 at dpi 10). *E. falciformis* isolate Brandenburg88 was more lethal for the *M. m. musculus* mice strains than for the other strains (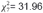, P<0.001; Table 2).

**Figure 2.**
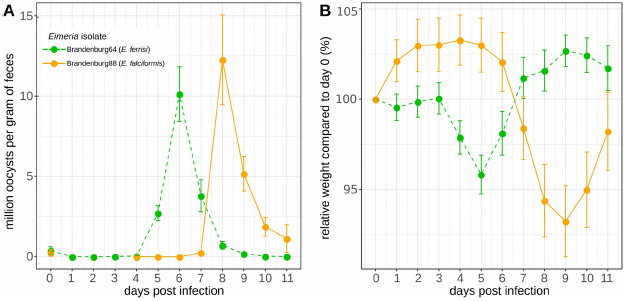
Parasite density (A) and host relative weight (B) during *Eimeria* infection. Parasite density is calculated as number of oocysts detected (in millions) per gram of feces, host relative weight is calculated as the percentage of weight compared to day 0. Mean and 95% CI are plotted for each parasite isolate. All mouse groups are pooled for each parasite isolate.

**Table 2.** Contingency table: number of mice and status at dpi 11 for each mouse group upon infection with E. falciformis isolate Brandenburg88.

| Mouse |  | status at dpi 11 |  |
| --- | --- | --- | --- |
| subspecies | group | alive | dead |
| Mmd | SCHUNT | 6 | 0 |
| Mmd | STRA | 7 | 0 |
| Mmd | SCHUNTxSTRA | 8 | 0 |
| Mmd-Mmm | STRAxBUSNA | 8 | 0 |
| Mmd-Mmm | SCHUNTxPWD | 6 | 0 |
| Mmm | PWDxBUSNA | 4 | 3 |
| Mmm | BUSNA | 3 | 4 |
| Mmm | PWD | 1 | 6 |
| total |  | 43 | 13 |

### 2. Comparison of resistance-tolerance coupling between *E. ferrisi* and *E. falciformis*

#### 2.1. Differences in resistance and tolerance between mouse groups depends on the parasite

Considering all mice infected with either *E. ferrisi* isolate Brandenburg 64 and *E. falciformis* isolate Brandenburg 88, we found our proxy for (inverse of) resistance (maximum number of OPG) to be statistically different between mouse groups, parasite isolates and their interaction (LRT: mouse groups: G=55.5, df=28, P<0.01; parasite isolates: G=40.5, df=16, P<0.001; interaction: G=27.9, df=14, P=0.015). Results were similar for our proxy for tolerance (LRT: mouse groups: G=28.4, df=14, P=0.01; parasite isolates: G=20.1 df=8, P=0.01; interaction: G=18.8, df=7, P<0.01). Our proxy for impact on weight (maximum relative weight loss) was significantly different between mouse groups and parasite isolates, but not for their interaction (LRT: mouse groups: G=44.9, df=14, P<0.001; parasite isolates: G=33, df=8, P<0.001; interaction: G=7.5, df=7, P=0.38). For the latter model, impact on weight, post-hoc tests showed that the only statistical differences between two mouse groups within a parasite infection were found in *E. falciformis* infection, between PWD and STRA (Tukey HSD test, p-value = 0.02), PWD and STRAxBUSNA (Tukey HSD test, p-value = 0.03) and PWD and SCHUNTxPWD (Tukey HSD test, p-value = 0.02). No difference was found within one mouse group between the two parasite isolates at the 0.05 significance threshold.

We found that the mean predicted number of OPG varies with the mean predicted relative weight loss (LRT: G=10, df=2, P<0.01), differs between both parasites (LRT: G=8.9, df=2, P=0.012), and more importantly we found a significant interaction term (LRT: G=8.3, df=1, P<0.01). This means that the relationship between mean health-effect and mean resistance differs between the two *Eimeria* species infections. Then, we performed a linear regression of the mean predicted tolerance for each eight mouse groups as a response of the mean predicted OPG, parasite isolate and their interaction. In this case we found that the mean number of OPG varies along with tolerance (LRT: G=8.5, df=2, P=0.01) but does not statistically differ between both parasites (LRT: G=1.1, df=2, P=0.57), and the interaction term was not found significant (LRT: G=0.03, df=1, P=0.86). In this respect, the correlation between resistance and tolerance was not found to significantly differ between both parasites. Following these results, we looked at the coupling of resistance and tolerance within each of the two isolates.

#### 2.2. Resistance and tolerance to *E. ferrisi* isolate Brandenburg64 are uncoupled

We tested coupling between resistance and tolerance for *E. ferrisi* isolate Brandenburg64 in our eight mouse groups. First, we tested whether our proxies for resistance and tolerance were different between the mouse groups. We found the maximum number of OPG to be statistically different between mouse groups (LRT: G=26.6, df=7, P<0.001; Figure 3A). Tolerance was not found to significantly differ between mouse groups for this parasite isolate (LRT: G=6.8, df=7, P=0.45; Figure 3B).

**Figure 3.**
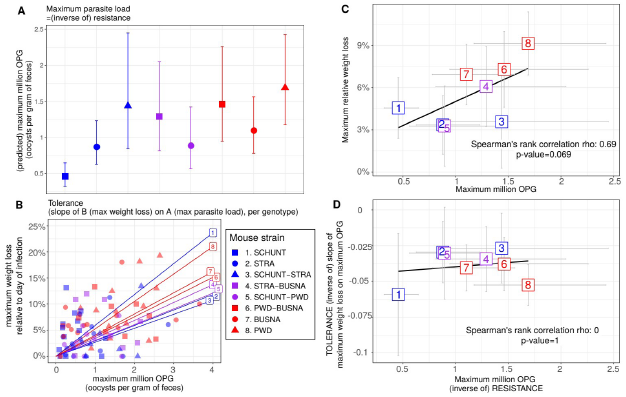
No indication of resistance-tolerance coupling for *E. ferrisi* isolate Brandenburg64. Colors represent mouse subspecies (blue: *M. m. domesticus*, red: *M. m. musculus*, purple: Mmd-Mmm). Left side: comparison of maximum oocysts per gram of feces used as a proxy for (inverse of) resistance (A) and tolerance (B) between mouse groups estimated by the slope of the linear regression with null intercept modelling maximum relative weight loss as a response of maximum oocysts per gram of feces, a steep slope corresponding to a low tolerance. Maximum number of OPG differs between mouse groups, but tolerance is similar. Right side: non significant positive correlation between mean maximum oocysts per gram of feces and mean relative weight loss (C) and absence of correlation between maximum oocysts per gram of feces used as a proxy for (inverse of) resistance and tolerance (D); Grey error bars represent 95% confidence intervals. Our results do not support coupling between resistance and tolerance *E. ferrisi* isolate Brandenburg64.

We found a non significant positive correlation between resistance (inverse of maximum number of OPG) and impact on health (maximum weight loss) (Spearman’s *ρ*=0.69, P=0.07, N=8; Figure 3C). Moreover, we did not find a correlation between resistance (inverse of maximum number of OPG) and tolerance (inverse of slope of maximum weight loss on maximum OPG) (Spearman’s *ρ*=0, P=1, N=8; Figure 3D).

In conclusion, we did not find indications of resistance-tolerance coupling for *E. ferrisi* isolate Brandenburg64, the different mouse groups infected by this parasite presenting a similar level of tolerance while showing an effect of quantitative resistance on health.

#### 2.3. Coupling between resistance and tolerance to *E. falciformis*

We then tested coupling between resistance and tolerance for *E. falciformis* isolate Brandenburg88 in our eight mouse groups. First, we tested if our proxies for resistance and tolerance were different between the mouse groups. We found the maximum number of OPG to be statistically different between mouse groups (LRT: G=28.6, df=14, P=0.012; Figure 4A). Contrary to our results on *E. ferrisi* isolate Brandenburg64, the tolerance slopes for *E. falciformis* isolate Brandenburg88 were different between mouse groups (LRT: G=13.9, df=7, P=0.05; Figure 4B).

**Figure 4.**
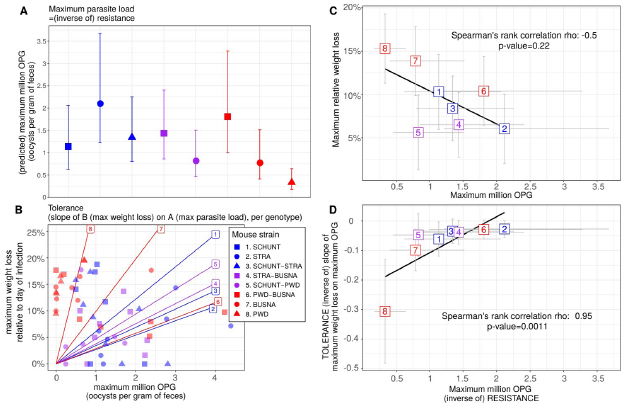
Coupling between resistance and tolerance for *E. falciformis* isolate Brandenburg88. Colors represent mouse subspecies (blue: *M. m. domesticus*, red: *M. m. musculus*, purple: Mmd-Mmm). Left side: comparison of maximum oocysts per gram of feces used as a proxy for (inverse of) resistance (A) and tolerance between mouse groups estimated by the slope of the linear regression with null intercept modelling maximum relative weight loss as a response of maximum oocysts per gram of feces, a steep slope corresponding to a low tolerance (B). Maximum number of OPG and tolerance differ between mouse groups. Right side: non significant negative correlation between mean maximum oocysts per gram of feces and mean relative weight loss (C) and strong positive correlation between maximum oocysts per gram of feces used as a proxy for inverse of resistance and tolerance (corresponding to a negative correlation between resistance and tolerance) (D); Grey error bars represent 95% confidence intervals. Our results support coupling between resistance and tolerance *E. falciformis* isolate Brandenburg88.

We detected a strong negative correlation between (inverse of) resistance (maximum number of OPG) and tolerance (inverse of slope of maximum weight loss on maximum OPG) (Spearman’s *ρ*=-0.95, P=0.001; Figure 4D). This result was robust to the exclusion of the extreme point corresponding to mouse strain PWD (point 8 in Figure 4D; Spearman’s *ρ*=-0.93, P<0.01).

We conclude that this correlation is unlikely a statistical artifact, as (1) mouse groups present statistically different values of resistance and tolerance (see 2.1) and (2) we found a (non significant) negative correlation between resistance (inverse of maximum number of OPG) and impact on health (maximum weight loss) (Spearman’s *ρ*=-0.5, P=0.22; Figure 4C), indicating that mouse groups losing more weight also shed less parasites.

We conclude that our results indicate the presence of negative resistance-tolerance coupling for *E. falciformis* isolate Brandenburg88.

## Discussion

In this study, we assessed resistance and tolerance to two closely related parasites, *E. ferrisi* and *E. falciformis*, in four mouse strains and their intra- and intersubspecific hybrids. Understanding this coupling has two major implications:

From a practical “measurement” perspective we can ask whether tolerance can be predicted from resistance, as the latter is easier to measure (e.g. in field sampling). Many studies assess the impact of parasites on host fitness based on resistance. If, as we found in the present study, resistance and tolerance are decoupled this can be misleading. In our host system, the house mice, for example, it has been shown that hybrids between *M. m. domesticus* and *M. m. musculus* are more resistant to parasites (Baird et al. 2012; Balard et al. 2020), including *Eimeria*, but tolerance could not be measured under natural conditions (Balard et al. 2020). The effect of parasites on host fitness in the evolution of the house mouse hybrid zone is thus still rather ambiguous (Baird and Goüy de Bellocq 2019). We show that careful distinction between parasite species is necessary when analysing parasite host interaction (see also Jarquín-Díaz et al. 2019) and that it is indispensable to measure both resistance and tolerance in *Eimeria* infections of house mice.

In this work we used the concept of tolerance as used originally in the plant literature and later on transferred to animal studies (Fineblum and Rausher 1995). This concept of tolerance can be criticised, as it links tolerance mathematically to resistance. Nevertheless, we argue that this view is biologically meaningful considering resistance and tolerance as a double defence system, one step limiting the parasite multiplication, the other limiting the impact of this multiplication on fitness-related traits. To limit the possibility of statistical artifact, our approach did not only consist in calculating correlations between resistance and tolerance, but also in testing differences in resistance, impact on health and tolerance. Of note, a positive correlation between mean health-effect and mean resistance of each host strains could indicate some host strains having few parasites-few effects on health, and others more parasites-more effects on health; This configuration would limit the possibility of detecting an actual resistance-tolerance trade-off by lack of a full range of resistance values. For this reason, our approach consisted in testing the “coupling” between resistance and tolerance, that is (1) a genotype-level correlation between tolerance and resistance additionally supported by (2) the absence of positive correlation between health-effect and resistance. We argue that this additional step increases the confidence in the presence of a biologically meaningful negative correlation between resistance and tolerance, likely implying a trade-off.

Differences between parasite species could explain the evolution of different strategies: *E. ferrisi* commits to sexual reproduction after a relatively short time with few cycles of asexual expansion (Al-khlifeh et al. 2019; Ankrom, Chobotar, and Ernst 1975), while *E. falciformis* has a relatively longer life cycle (Al-khlifeh et al. 2019; Haberkorn 1970). As *E. ferrisi* infections do not reach extremely high intensities, high tolerance might be the optimal strategy for both house mouse subspecies. Resistance could then evolve relatively freely without any major impact of the parasite on the hosts’ health. In the case of *E. falciformis*, the long life cycle might lead to high tissue load. Tissue damage is observed during sexual reproduction for this parasite (Ehret et al. 2017) and might mean that a certain level of resistance is required. On the other hand, immunopathology has been observed in advanced *E. falciformis* infections (Stange et al. 2012). These intrinsic characteristics of *E. falciformis* might lead to multiple different optima for resistance and tolerance, leading to a trade-off.

More generally, from an evolutionary perspective, coupling between resistance and tolerance might help determine if coevolution between host and parasite can be expected: a host-parasite system in which one finds negative coupling between tolerance and resistance would be an especially promising system for studies of host-parasite coevolution. Indeed, coevolution in host-parasite systems is often assumed but rarely proven (Woolhouse et al. 2002). Janzen (1980) notes that not all parasite-host systems are coevolving. The presence of efficient host defences against a given parasite is not necessarily produced in response to this parasite specifically and the parasite does not necessarily respond specifically. In the mouse-*E. ferrisi* system, where resistance and tolerance are decoupled, host and parasite fitness might be decoupled as a result, making host-parasite coevolution less likely. In the mouse-*E. falciformis* system we found a negative coupling between tolerance and resistance, making coevolution between host and parasite more likely.

In conclusion, we show that the coupling between resistance and tolerance can differ between closely related parasite species and we argue that this trait of a host-parasite system determines the questions to be best approached with a particular parasite.

## Authors contributions

AB, JP and EH designed the experiment and analysis. LD and JP provided the research material. AB, VHJD, JJ, VM and FB carried out the experiment. AB performed the analysis. AB and EH wrote the manuscript, with major contribution from JP and feedback from all the authors. Funding: This work was funded by the German Research Foundation (DFG) Grant [HE 7320/1-1] to EH. VHJ is an associated student of GRK 2046 funded by the DFG. The maintenance of wild-derived strains was supported by the ROSE program from Czech Academy of Sciences and the Czech Science Foundation (project 16-23773S) to JP.

## Data Accessibility

-Code and full data: Zenodo doi: 10.5281/zenodo.3911935

## Authors Contributions

AB, JP and EH designed the experiment and analysis. LD and JP provided the research material. AB, VHJD, JJ, VM and FB carried out the experiment. AB performed the analysis. AB and EH wrote the manuscript, with major contribution from JP and feedback from all the authors.

## Acknowledgements

This work was funded by the German Research Foundation (DFG) Grant [HE 7320/1-1] to EH. VHJ is an associated student of GRK 2046 funded by the DFG. The maintenance of wild-derived strains was supported by the CAS within the program of the Strategy AV 21 and the Czech Science Foundation (project 16-23773S) to JP.

## Appendix

**Appendix 1.**
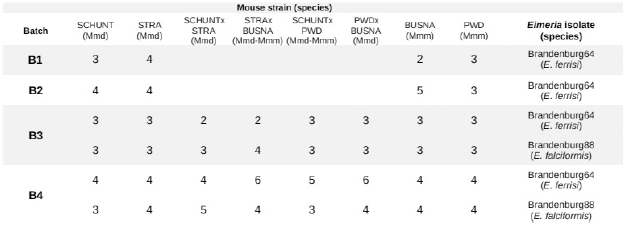
Chronology of experimental infections.

**Appendix 2.**
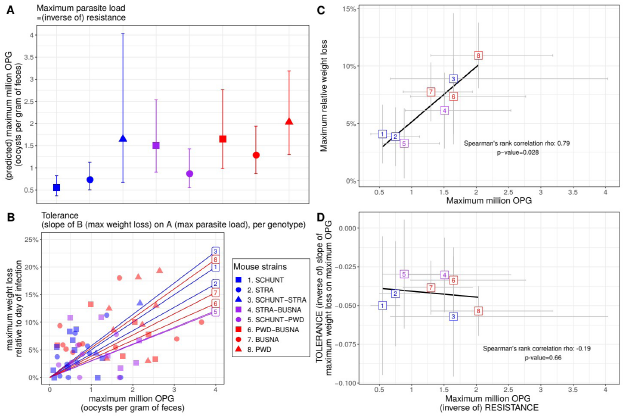
No indication of resistance-tolerance coupling for *E. ferrisi* isolate Brandenburg64 in the conservative dataset. Colors represent mouse subspecies (blue: *M. m. domesticus*, red: *M. m. musculus*, purple: Mmd-Mmm). Left side: comparison of maximum oocysts per gram of feces used as a proxy for (inverse of) resistance (A) and tolerance (B) between mouse groups estimated by the slope of the linear regression with null intercept modelling maximum relative weight loss as a response of maximum oocysts per gram of feces, a steep slope corresponding to a low tolerance. Maximum number of OPG differs between mouse groups, but tolerance is similar. Right side: positive correlation between mean maximum oocysts per gram of feces and mean relative weight loss (C) and absence of correlation between maximum oocysts per gram of feces used as a proxy for (inverse of) resistance and tolerance (D); Grey error bars represent 95% confidence intervals. Our results do not support coupling between resistance and tolerance *E. ferrisi* isolate Brandenburg64.

**Appendix 3.**
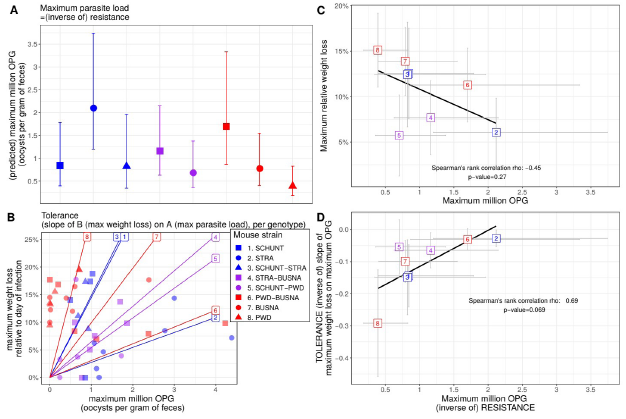
Coupling between resistance and tolerance for *E. falciformis* isolate Brandenburg88 in the conservative dataset. Colors represent mouse subspecies (blue: *M. m. domesticus*, red: *M. m. musculus*, purple: Mmd-Mmm). Left side: comparison of maximum oocysts per gram of feces used as a proxy for (inverse of) resistance (A) and tolerance (B) between mouse groups estimated by the slope of the linear regression with null intercept modelling maximum relative weight loss as a response of maximum oocysts per gram of feces, a steep slope corresponding to a low tolerance. Maximum number of OPG and tolerance differ between mouse groups. Right side: non significant negative correlation between mean maximum oocysts per gram of feces and mean relative weight loss (C) and strong positive correlation between maximum oocysts per gram of feces used as a proxy for inverse of resistance and tolerance (corresponding to a negative correlation between resistance and tolerance) (D); Grey error bars represent 95% confidence intervals. Our results support coupling between resistance and tolerance *E. falciformis* isolate Brandenburg88.

