## Supplementary Table S1 for "Coupling between tolerance and resistance for two related *Eimeria* parasite species" (from another version of this preprint: version 2, 2020-06-27)

**Supplementary Table S1. Chronology of experimental infections.**

| | Mouse strain (species) | | | | | | | | |
|---|---|---|---|---|---|---|---|---|---|
| **Batch** | SCHUNT (Mmd) | STRA (Mmd) | SCHUNTx STRA (Mmd) | STRAx BUSNA (Mmd-Mmm) | SCHUNTx PWD (Mmd-Mmm) | PWDx BUSNA (Mmd) | BUSNA (Mmm) | PWD (Mmm) | ***Eimeria* isolate (species)** |
| **B1** | 2 | 3 | | | | | 2 | 3 | Brandenburg139 (*E. ferrisi*) |
| | 3 | 4 | | | | | 2 | 3 | Brandenburg64 (*E. ferrisi*) |
| **B2** | 5 | 3 | | | | | 4 | 3 | Brandenburg139 (*E. ferrisi*) |
| | 4 | 4 | | | | | 5 | 3 | Brandenburg64 (*E. ferrisi*) |
| **B3** | 3 | 3 | 2 | 2 | 3 | 3 | 3 | 3 | Brandenburg64 (*E. ferrisi*) |
| | 3 | 3 | 3 | 4 | 3 | 3 | 3 | 3 | Brandenburg88 (*E. falciformis*) |
| **B4** | 4 | 4 | 4 | 6 | 5 | 6 | 4 | 4 | Brandenburg64 (*E. ferrisi*) |
| | 3 | 4 | 5 | 4 | 3 | 4 | 4 | 4 | Brandenburg88 (*E. falciformis*) |
