## Supplementary Material S2 for "Coupling between tolerance and resistance for two related *Eimeria* parasite species" (from another version of this preprint: version 2, 2020-06-27)

### Supplementary material S2. Conservative dataset (N=118)

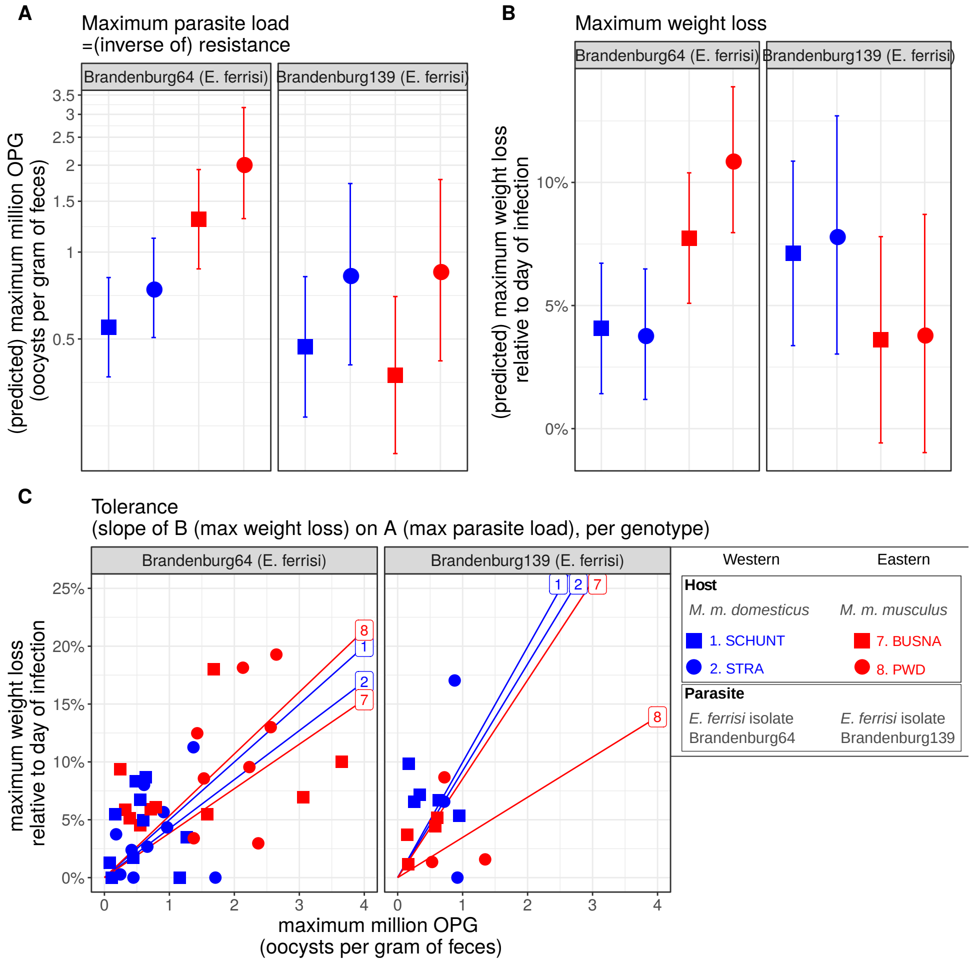


**Figure S2.1. Comparison of resistance, impact on weight and tolerance between mouse strains for both *Eimeria ferrisi* isolates.** (A) Maximum oocysts per gram of feces used as a proxy for (inverse of) resistance; (B) Impact on host health measured as the maximum weight loss during patent period relative to starting weight (%); (C) Tolerance estimated by the slope of the linear regression with null intercept modelling maximum relative weight loss as a response of maximum oocysts per gram of feces. A steep slope corresponds to a low tolerance. We did not detect (A) either higher parasite shedding of the Eastern parasite isolate in Eastern mouse strains and *vice versa* (LRT interaction factor mouse strain-parasite isolate: G=6.9, df=3, P=0.74) or (C) higher tolerance of Eastern hosts infected by Eastern parasite isolate and *vice versa* (LRT interaction factor mouse strain-parasite isolate: G=3.1, df=3, p=0.38), thus our results do not support the hypothesis of local adaptation between *E. ferrisi* and its host.

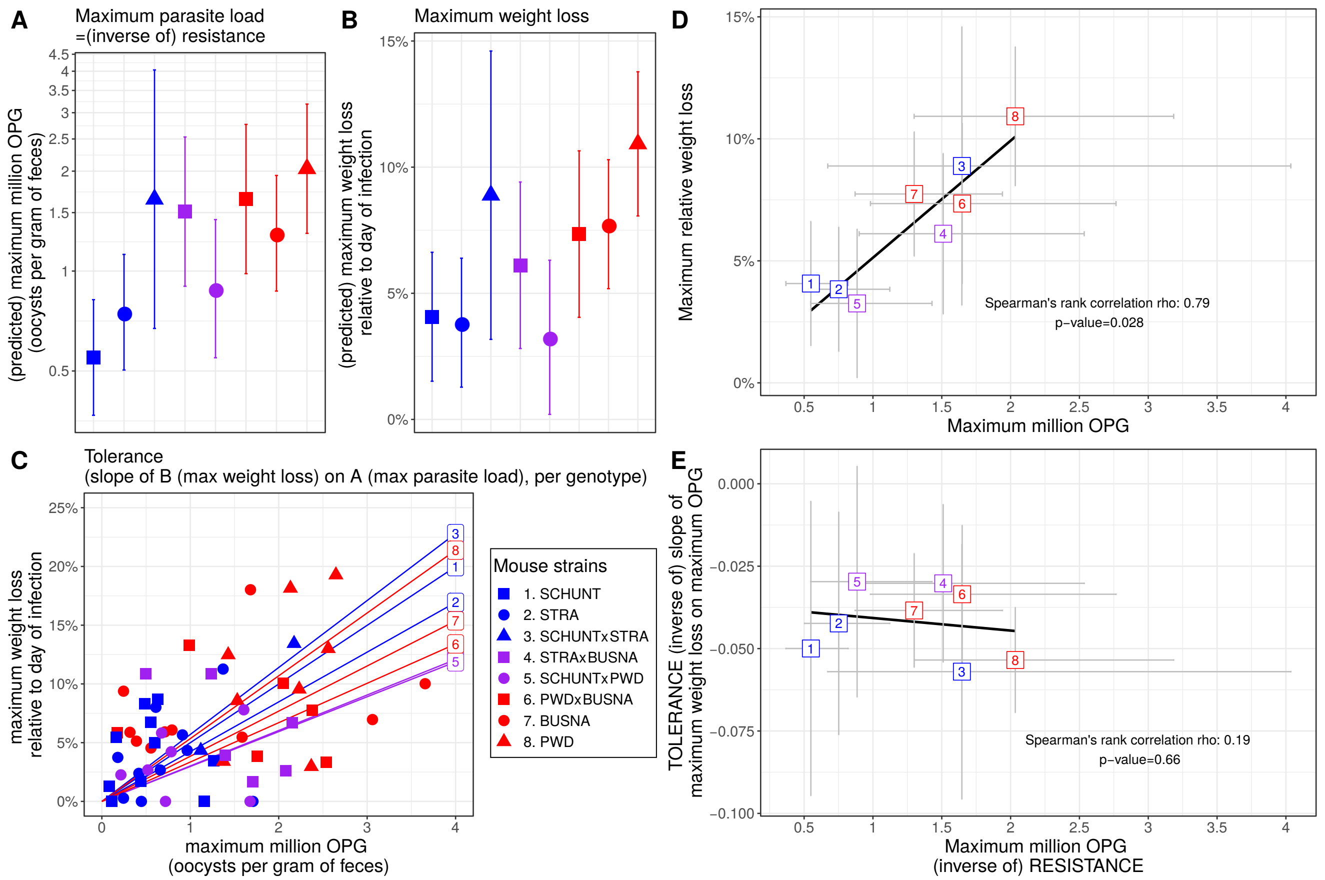


**Figure S2.2. No indication of resistance-tolerance coupling for *E. ferrisi* isolate Brandenburg64.** Colors represent mouse subspecies (blue: Mmd, red: Mmm, purple: Mmd-Mmm). Left side: comparison of maximum oocysts per gram of feces used as a proxy for (inverse of) resistance (A), impact on weight measured as the maximum weight loss during patent period relative to starting weight (B) and tolerance between mouse groups estimated by the slope of the linear regression with null intercept modelling maximum relative weight loss as a response of maximum oocysts per gram of feces, a steep slope corresponding to a low tolerance (C). Maximum number of OPG and relative weight loss differ between mouse groups (LRT: maximum number of OPG: G=22.6, df=7, p=0.002; maximum relative weight loss: G=21.7, df=7, p=0.0028), but tolerance is similar (LRT: G=5.4, df=7, p=0.62). Right side: non significant positive correlation between mean maximum oocysts per gram of feces and mean relative weight loss (D) and absence of correlation between maximum oocysts per gram of feces used as a proxy for (inverse of) resistance and tolerance (E); Grey error bars represent 95% confidence intervals. Our results do not support coupling between resistance and tolerance *E. ferrisi* isolate Brandenburg64.

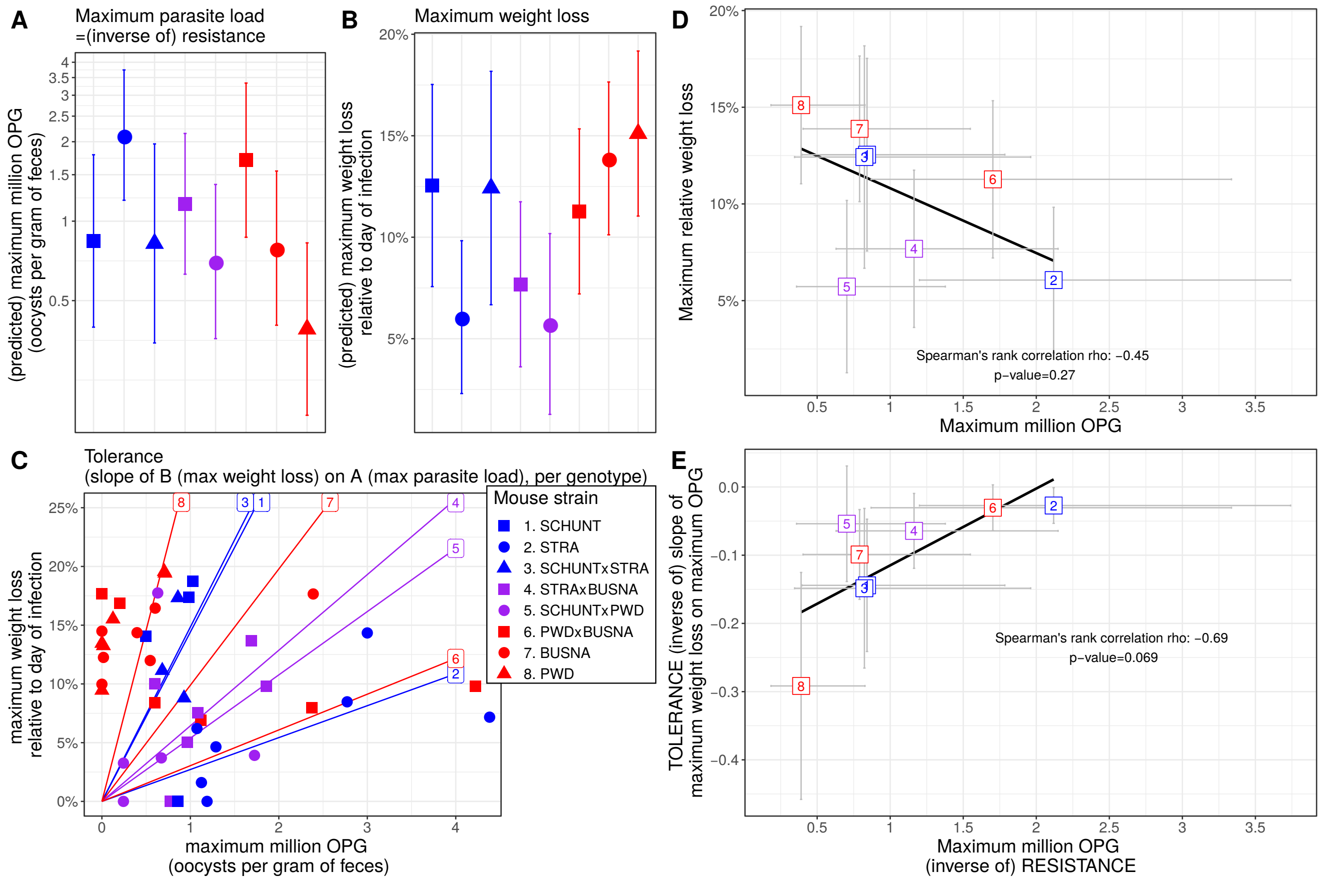


**Figure S2.3. Coupling between resistance and tolerance for *E. falciformis* isolate Brandenburg88**. Colors represent mouse subspecies (blue: Mmd, red: Mmm, purple: Mmd-Mmm). Left side: comparison of maximum oocysts per gram of feces used as a proxy for (inverse of) resistance (A), impact on weight measured as the maximum weight loss during patent period relative to starting weight (B) and tolerance between mouse groups estimated by the slope of the linear regression with null intercept modelling maximum relative weight loss as a response of maximum oocysts per gram of feces, a steep slope corresponding to a low tolerance (C). Maximum number of OPG, relative weight loss and tolerance differ between mouse groups (LRT: maximum number of OPG: G=24, df=14, p=0.046; maximum relative weight loss: G=20.1, df=7, p=0.005; tolerance: G=20.2, df=7, p=0.0051). Right side: non significant negative correlation between mean maximum oocysts per gram of feces and mean relative weight loss (D) and non significant negative correlation between maximum oocysts per gram of feces used as a proxy for (inverse of) resistance and tolerance (E); Grey error bars represent 95% confidence intervals. Our results present indications of coupling between resistance and tolerance *E. falciformis* isolate Brandenburg88, with lower support than the full dataset likely due to the lower statistical power.
